# The multiplicity of infection does not affect interactions between *Cryptococcus neoformans* and macrophages

**DOI:** 10.1101/2020.02.05.936427

**Authors:** Katherine Pline, Simon Johnston

**Affiliations:** Department of Infection, Immunity and Cardiovascular Disease, Bateson Centre and Florey Institute, University of Sheffield, UK

## Abstract

Major determinants of the outcome of infection include growth of the pathogen and response of immune cells such as macrophages. *Cryptococcus neoformans* is a fungal pathogen which may grow within the extracellular environment, or may exploit host macrophages as a niche for replication and dissemination. The clinical outcome of cryptococcal infection varies widely between individuals even when key host and pathogen molecular factors are the same. For a broad range of infections altering pathogen density is known to influence progression and outcome of infection by affecting immune response and pathogen biology (e.g. via innate immune signalling or microbial quorum sensing). Here, using time lapse imaging of murine cell line and human primary macrophages *in vitro*, we examined the effect of altering pathogen density on the interactions of macrophages with cryptococci. We find that increasing fungal burden over several orders of magnitude did not increase or decrease phagocytosis by murine J774 macrophage-like cells or human monocyte-derived macrophages, illustrating neither dose-dependent immune activation nor dampening of the phagocytic response. Furthermore, increasing fungal density alone was not sufficient to alter the ability of cryptococci to grow intracellularly and has no significant effect on fungal doubling time for cryptococci that were intracellular or extracellular. This suggested that individual macrophage-*Cryptococcus* interactions were not affected by even large changes in fungal density, an important finding in understanding what determines the outcome of cryptococcal infection.

## Introduction

During infection, pathogen growth and immune response largely determine the progression and outcome of infection. Pathogen replication and dissemination drive infection progression, while the immune response attempts to control the growth, spread, and ultimately sterilise the infecting organism. Increasing infection burden eventually overwhelms the actions of the immune system and leads to uncontrolled pathogen growth. However, especially early in infection, increasing pathogen density may also induce a more robust and definitive immune response that leads to infection clearance. Therefore, the effect of pathogen density on infection growth and immune response must be carefully considered to understand infection dynamics.

For pathogens, quorum sensing (QS) is a density-dependent coordinated response amongst microbes to modulate gene expression and phenotype at threshold signalling molecule concentrations in their environment within a host during infection. For example, in culture when a high population density is achieved, *Candida albicans* produces a signalling molecule at sufficiently high concentrations which shortens time the pathogen spends in the lag phase of growth (Chen et al., 2004); thus, pathogenic density and subsequent signalling molecule concentration enhances the growth of this pathogen, which may directly impact the progression and control of infection. Pathogens may also respond to population density by coordinating survival mechanisms, morphology, and the expression of virulence factors. Because this density-dependent response may enhance the pathogenicity of invading pathogens, it is important to understand if pathogens coordinate growth and behaviour based on population density, and to determine what effect this may have on immune response and the progression of infection.

Professional phagocytes such as macrophages are key cells in the cell mediated innate immune response, as well as coordinators and effector cells in the wider immune response. The phagocytosis of pathogens by macrophages early in infection is key in controlling the growth and spread of infection. Therefore, with respect to the immune response, the effect of increasing pathogenic density on phagocytic uptake and control must also be considered. For example, at high density the fungal pathogen *Histoplasma capsulatum* produces cell wall polysaccharides which may prevent recognition by host phagocytes, inhibiting uptake and destruction by macrophages (Rappleye et al., 2007). Furthermore, because these polysaccharides are produced by the yeast while replicating within host macrophages where intraphagosomal concentration of QS signalling molecules is likely to be high, it may be a potential mechanism by which the pathogen uses quorum sensing to detect when it is in a phagosome (Kügler et al., 2000). This signalling may induce the expression of virulence factors by the yeasts and result in the enhanced parasitism of the host macrophage. Because density-dependent responses by pathogens may not only impair uptake by phagocytes, but may enable enhanced intracellular parasitism of host cells, the effect of pathogen density on phagocyte response must be investigated to determine how this affects infection outcome.

*Cryptococcus neoformans* is a pathogenic yeast which may be phagocytosed and destroyed by host macrophages, or may colonise the macrophage intracellular niche to replicate intracellularly and disseminate infection (Johnston and May, 2013; Rudman et al., 2019). As such, interactions between this important host phagocyte and the invading fungal pathogen may have the potential to affect the variable infection outcomes seen in human cryptococcosis (Lee et al., 2011; Pappas et al., 2001). Though QS has been shown to be important to the pathogenicity of many fungal pathogens, density-dependent cryptococcal behaviour remains poorly understood. Evidence for cryptococcal QS comes from the observation that while the *Δtup1* mutant strain of *C. neoformans* failed to grow at a culture density of less than 1,000 cells, conditioned media from high-density cultures induced fungal growth at this low density (Lee et al., 2007). The peptide CQS1 was isolated and attributed with inducing a density-dependent QS response (Lee et al., 2007). However, while there is evidence supporting the idea that *C. neoformans* may exhibit density-dependent growth and phenotypic responses in culture, such effects with respect to interactions with macrophages remains largely unexplored.

Here, using time lapse analysis of murine macrophage-like cells and human monocyte-derived macrophages, we show that increasing cryptococcal multiplicity of infection does not affect phagocytosis by macrophages or fungal replication. Furthermore, we show that uptake of cryptococci by macrophages increases in proportion with initial infection dose, and provide evidence that each macrophage-*Cryptococcus* encounter should be considered an independent infection. Thus, cryptococcal replication and macrophage responses are neither enhanced nor inhibited by increasing multiplicity of infection (MOI), and fungal density-dependent signalling does not appear to affect host-pathogen interactions during *in vitro* cryptococcal infection.

## Results

### There is no activation or inhibition of phagocytosis of *C. neoformans* by J774 macrophages dependent on fungal density

We investigated how increasing cryptococcal MOI affected uptake by J774 murine macrophage-like cells (Voelz et al., 2009). 10^5^ activated J774 macrophages were infected with 10^2^-10^5^ opsonised KN99 GFP *C. neoformans* (an MOI of 0.001-1) (Gibson et al., 2018). This timepoint was 0 hours, and infection was followed over 18 hours (Figure 1a). We then calculated the proportion of the total macrophages which contained intracellular cryptococci over this increasing MOI. We reasoned that if increasing fungal burden overwhelmed the macrophages’ ability to phagocytose cryptococci, as fungal burden increased the proportion of macrophages that contained cryptococci would plateau or decrease. Alternatively, if the proportion of macrophages containing intracellular cryptococci increased disproportionately to the level of fungal infection, this might indicate enhanced phagocytic activation of macrophages. Time lapse analysis showed there was a linear relationship between fungal burden and the percentage of macrophages with intracellular cryptococci, both at 0 and 18 hours post infection (Figures 1b and c; linear regression for 0h p < 0.0001, R^2^ = 0.892, for 18h p < 0.0001, R^2^ = 0.859). The percentage of infected macrophages was not significantly different between 0 and 18 hours (Figure 1d Mann-Whitney test p = 0.700), and macrophage proliferation was not significantly affected by increasing fungal burden (Figure 1e Kruskal-Wallis test p = 0.0788). These results illustrate that J774 macrophages showed neither enhanced nor reduced phagocytosis over three orders of magnitude of cryptococcal infection, with phagocytosis increasing linearly with infection burden. Similar results were achieved through flow cytometry analysis (Supplemental figures 1a-f; Voelz et al. 2011).

**Fig. 1.**
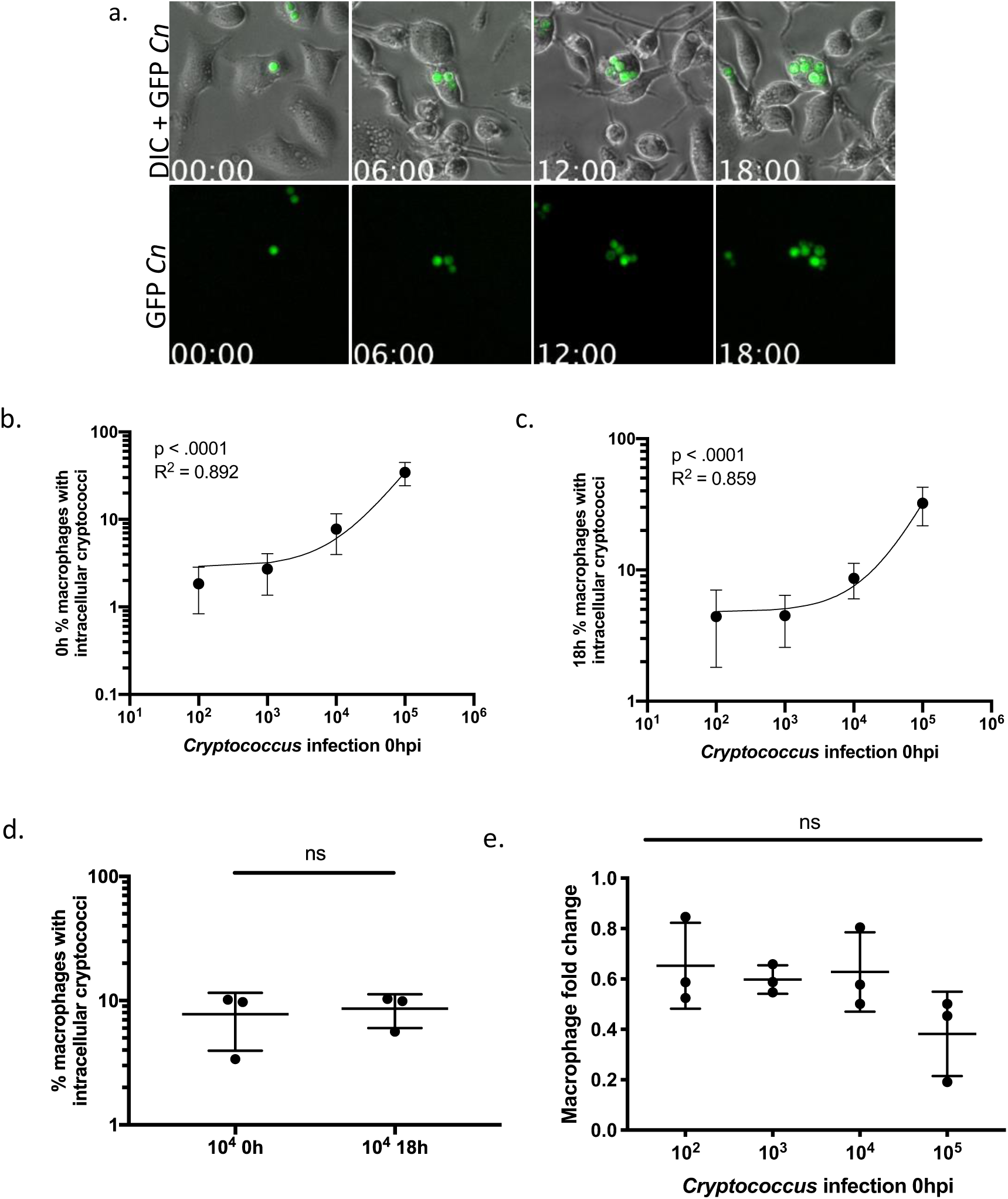
Phagocytosis of KN99 GFP by J774 macrophages. J774 macrophages were infected with KN99 GFP *C. neoformans* (a), infections were followed for 18 hours and the percentage of macrophages containing intracellular cryptococci was quantified. At 0h (b) and 18h (c), there was a linear relationship between the infecting fungal burden and the percentage of macrophages with intracellular cryptococci (0h linear regression p < 0.0001, R^2^ = 0.892; 18h linear regression p < 0.0001, R^2^ = 0.859. There was no significant difference between the proportion of macrophages containing intracellular cryptococci at 0h and 18h (d, Mann-Whitney test p = 0.700). Only one initial fungal dose shown here for clarity, but results were similarly non-significant in each comparison. There was no significant effect of cryptococcal MOI on macrophage proliferation (e Kruskal-Wallis test p = 0.0788, no pairwise significance). Three repeats were performed, with no less than 250 macrophages considered per condition.

### The multiplicity of *C. neoformans* infection does not enhance or inhibit the ability of cryptococci to replicate intracellularly within J774 macrophages

As population density may affect the ability of pathogens to proliferate, and may therefore affect the progression of infection, we next measured how increasing fungal burden affected the replication of cryptococci. We reasoned that if cryptococci cooperated to enhance replication as fungal burden increased, as seen in many examples of quorum sensing, that the fold change relative to initial burden would increase with dose. Alternatively, if increasing burden induced cryptococci to slow growth, the relative fold change would decrease as fungal burden increased. With the aim of understanding how increasing MOI affects macrophage-*Cryptococcus* interactions, we quantified the intracellular cryptococcal fold change relative to initial fungal burden in the time lapse assay above. This revealed that increasing fungal burden did not enhance or inhibit the ability of intracellular cryptococci to replicate (Figure 2a; relative fold change one-way ANOVA p = 0.475). This suggests that increasing fungal burden did not increase or decrease the ability of cryptococci to replicate within host macrophages. Similar results were achieved by flow cytometry analysis (Supplemental figure 1f).

**Fig. 2.**
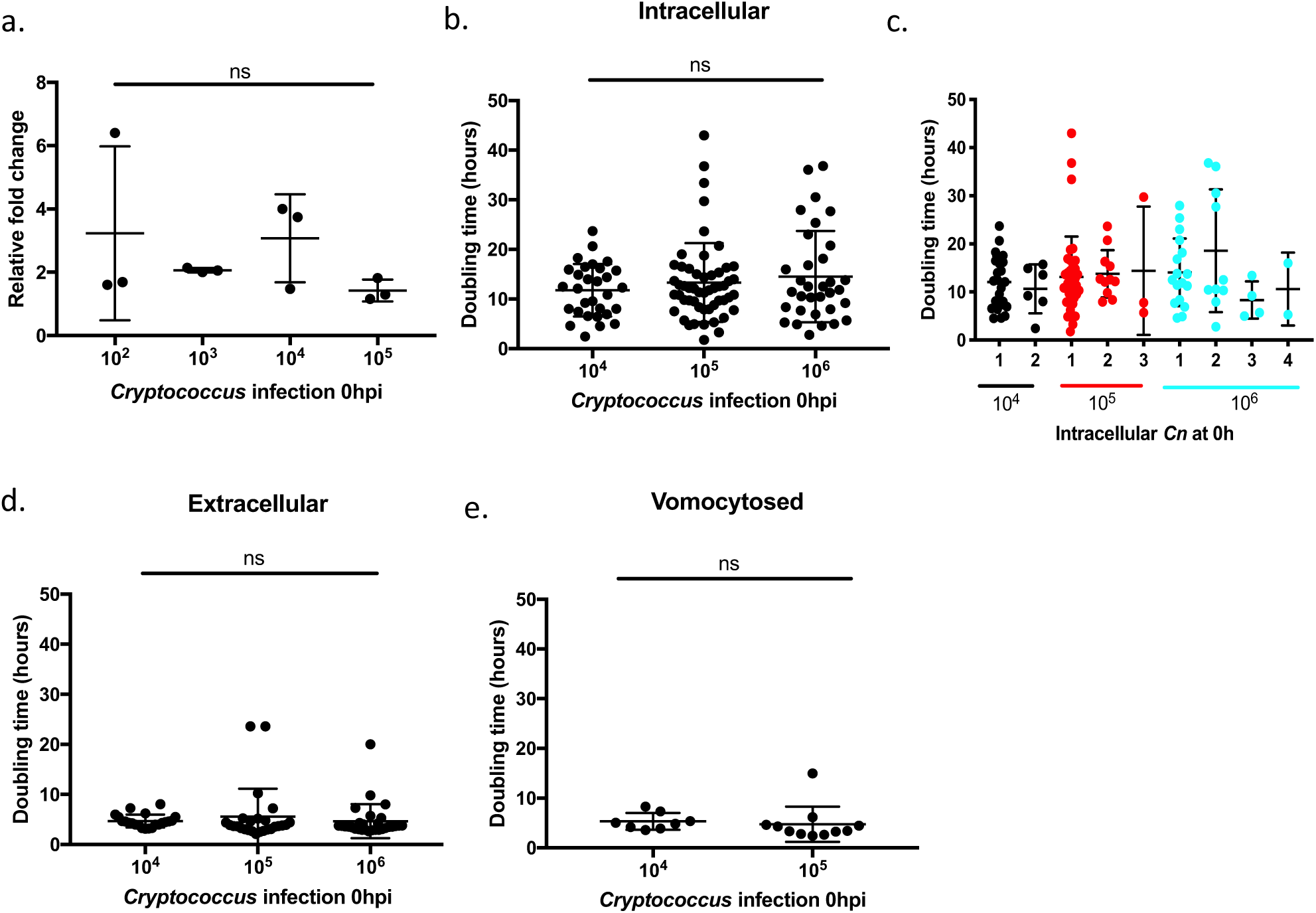
Cryptococcal doubling time during J774 macrophage infection. J774 macrophages were infected with an increasing inoculum of KN99 GFP *C. neoformans*, and individual cryptococcal cells were followed for up to 24 hours to quantify doubling time. There was no significant difference in relative fold change of intracellular cryptococcal cells after 18 hours of infection (a, one-way ANOVA p = 0.475). Doubling time of intracellular cryptococci was not affected by initial fungal dose (b, one-way ANOVA p = 0.3724, no pairwise significance), and stratification by the initial number of phagocytosed fungal cells per macrophage also showed no difference in doubling time (c, one-way ANOVA p = 0.5572, no pairwise significance). There was no difference in doubling time amongst extracellular (d, one-way ANOVA p = 0.6348), or vomocytosed cryptococci (e, only 2 levels of infection considered due to the difficulty in following vomocytosed cells late in infection at the highest fungal burden; Mann-Whitney test p = 0.6899). Three repeats were performed, with the average of three replicates per condition per repeat shown. Up to eleven cells were followed per replicate.

While calculating fungal fold change is a good indicator of the ability of *Cryptococcus* to proliferate at increasing levels of fungal burden, it does not provide a specific measurement of the rate of replication. Comparing intracellular and extracellular fungal replication rates provides a measure of macrophage control of infection. As such, we performed the same infection assay as above and quantified the growth of individual cryptococcal cells within macrophages, outside of macrophages, and after vomocytosis from macrophages. A higher MOI was used for this assay to produce a higher number of infected macrophages for time lapse imaging; 10^5^ macrophages were infected with 10^3^-10^6^ KN99 GFP *C. neoformans*, and infection was followed by time lapse analysis for 24 hours. Following cryptococcal replication within individual macrophages showed that as the initial fungal burden increased, the mean doubling time of individual cryptococci neither increased nor decreased (Figure 2b; one-way ANOVA p = 0.3724). This suggests that increasing MOI did not induce a faster rate of intracellular cryptococcal proliferation, and that macrophage control of intracellular replication was not affected by initial fungal burden. While there was no difference at a population level, we hypothesised that macrophages with a higher initial burden might have impaired control of cryptococcal growth. Therefore, we also quantified the doubling times of cryptococci within macrophages by the initial number of cryptococci within each macrophage (i.e. the number of cryptococci within the macrophage phagosome). However, as the intracellular cryptococcal burden increased, there was no significant difference in doubling time (Figure 2c one-way ANOVA p = 0.5572). These combined results show that as the level of fungal burden increases, cryptococci do not cooperate to enhance or reduce intracellular proliferation, and that macrophage control of intracellular replication is not affected by increasing burden.

As *C. neoformans* is a facultative intracellular pathogen which may replicate in the extracellular environment, and as macrophages may affect the extracellular growth of cryptococci by secreting antimicrobial effector molecules, we also measured how increasing fungal burden affected growth of cryptococci which were extracellular in the presence of macrophages. Following the replication individual cryptococci showed that increasing MOI did not affect the doubling time of extracellular cryptococci (Figure 2d one-way ANOVA p = 0.6348). Because cryptococci may exit non-lytically from macrophages in a process called vomocytosis, we also quantified the doubling time of cryptococci which vomocytosed from J774 macrophages. Again, there was no effect of infection burden on the doubling time of vomocytosed cryptococci (Figure 2e Mann-Whitney test p = 0.6899). These results show that over increasing fungal burdens, cryptococci did not cooperate to enhance replication rate. These results also indicate that macrophages did not exert enhanced or reduced control over extracellular cryptococcal proliferation, suggesting that increasing MOI did not induce the increased secretion antimicrobial factors by macrophages.

### There is no dose activation or inhibition of phagocytosis of cryptococci by human monocyte-derived macrophages

While J774 macrophages are a good model for *C. neoformans* phagocytosis and intracellular proliferation (Johnston and May, 2010; Ma et al., 2009; Voelz et al., 2009), there are differences between human cells and murine cells which means that the interaction between cryptococci and murine macrophages may not fully capture the nature of interactions between human macrophages and cryptococci (Ingersoll et al., 2010). We therefore chose to infect human monocyte-derived macrophages (MDMs) to see if human macrophages controlled cryptococcal infection in a similar manner to the murine macrophages. To determine if the effects of infection dose on macrophage phagocytosis and fungal replication observed in the presence of murine macrophages were similar for human cells, we performed the assays described above in the presence of human MDMs. We infected 10^5^ activated MDMs with 10^3^-10^6^ opsonised KN99 GFP *C. neoformans* for 2 hours, washed off extracellular cryptococci, and followed infection for 24 hours by time lapse imaging (Figure 3a). To discern if fungal dose affected the ability of human macrophages to phagocytose cryptococci, we quantified the proportion of macrophages with intracellular cryptococci over 18 hours of infection. The number of infected macrophages was only quantified at fungal burdens up to 10^5^ because phagocytosis was difficult to quantify at the highest dose.

**Fig. 3.**
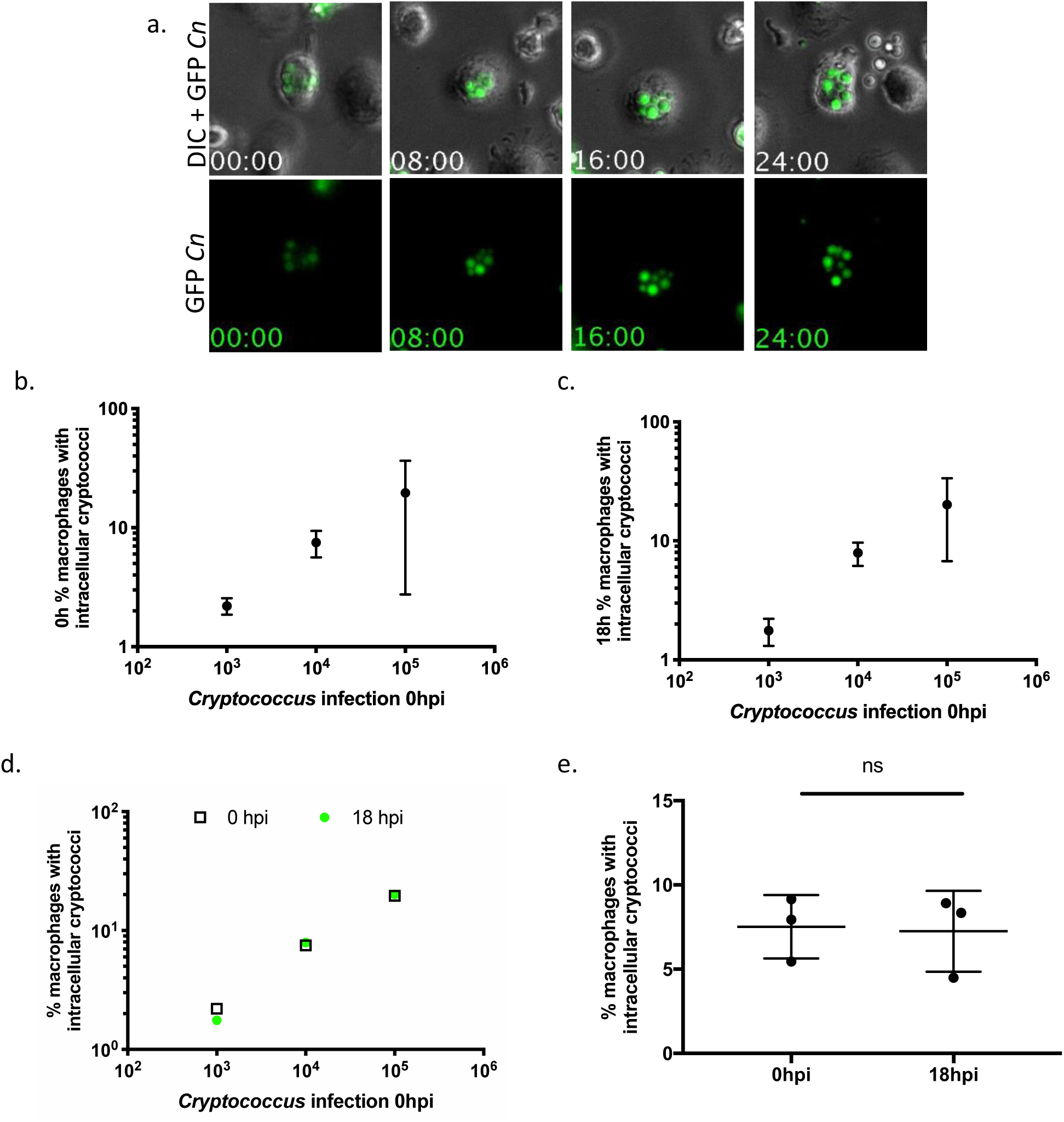
Phagocytosis of cryptococci by human monocyte derived macrophages. Human MDMs were infected for 24h with KN99 GFP *C. neoformans* (a). At the start of infection, the percentage of infected macrophages did not increase linearly with dose (b, linear regression p = 0.139). At 18h the percentage of infected macrophages did not increase linearly with initial fungal burden (c, linear regression p = 0.161). A comparison of the mean percentage of macrophages containing intracellular cryptococci at 0 and 18 hours (d, i.e. the means depicted in (a) and (b)). Transparent squares represent the mean percent of infected macrophages across repeats at 0 hpi, and green circles represent 18h. The mean percentage of macrophages with intracellular cryptococci increased with dose, though this increase was not linear (linear regression for 0h p = 0.139, for 18h p = 0.161). The proportion of macrophages which contained intracellular cryptococci did not change significantly after 18h (e, Mann-Whitney test p > 0.999, one dose shown for clarity no significant difference for each fungal dose). Three repeats were performed, with 2-3 replicates per initial fungal dose per repeat. The mean values for each dose within each repeat are shown.

At 0 hours, linear regression analysis showed that there was no significant linear relationship between initial fungal burden and the proportion of MDMs with intracellular cryptococci (Figure 3b linear regression p = 0.139). There was also no linear relationship between the initial fungal burden and the percentage of infected MDMs at 18h (Figure 3c linear regression p = 0.161). These results suggest that increasing the number of cryptococci within the environment potentially overwhelmed MDM phagocytic activity, as the percentage of macrophages with intracellular cryptococci at the highest fungal dose was less than what would be expected if there was a linear relationship between fungal burden and phagocytic uptake. However, it is likely that stochastic interactions between cryptococci and macrophages may have contributed to the high standard deviation in the percent of infected macrophages, failing to produce a linear relationship. Despite this, the mean proportion of infected macrophages did increase proportionally with initial fungal burden; considering only the mean percentage of macrophages infected without this variation shows that there was an increase in uptake with fungal dose, though this was not linear (Figure 3d; linear regression p = 0.139 for 0h, p = 0.161 for 18h). As with the J774 time lapse analysis, the percentage of macrophages with intracellular cryptococci did not change over 18 hours (Figure 3e; unpaired t-test p > 0.99, non-significant difference for each dose considered). These results suggest that though the percentage of infected macrophages increased with MOI, there may have been some inhibition of phagocytosis by MDMs at high infection burdens, suggesting that phagocytosis was affected by fungal dose *in vitro*; however, donor variation may have also contributed to this non-linear outcome.

### During human MDM infection, increasing cryptococcal burden did not affect fungal replication

As before, we hypothesised that if an increase in the number of cryptococci induced fungal cooperation and upregulation of replication, relative fold change would increase with MOI in MDMs. If increasing fungal dose resulted in a downregulation of replication, relative fold change would decrease with MOI. Time lapse analysis showed that as the initial fungal burden increased, there was no increase or decrease in the relative fungal fold change within human MDMs (Figure 4a one-way ANOVA p = 0.696). To determine if the level of cryptococcal infection had any effect on the replication rate of cryptococci, we followed individual intracellular and extracellular cryptococcal cells for up to 24 hours to allow doubling time to be determined. We then quantified the replication of individual cryptococci. Within MDMs, as the total initial fungal burden increased, there was no significant increase or decrease in doubling time of intracellular cryptococci (Figure 4b one-way ANOVA p = 0.264). This suggested that the presence of additional cryptococci did not result in signalling amongst the fungal population that induced individual fungal cells to replicate at a faster rate. Interestingly, as the number of cryptococci inside of separate macrophages increased (i.e. starting intracellular cryptococci within each macrophage), the doubling time of the intracellular cryptococci increased, irrespective of initial infection dose (Figure 4c). Finally, as initial fungal burden increased there was no effect of dose on the extracellular doubling rate (Figure 4d one-way ANOVA p = 0.844). This indicates that increasing fungal load did not induce extracellular cryptococci to regulate replication rate in a dose-dependent manner. Taken together these results show that as the initial fungal burden increased, the cryptococci may signal amongst one another to downregulate replication rate, but only within subpopulations inside of individual MDMs.

**Fig. 4.**
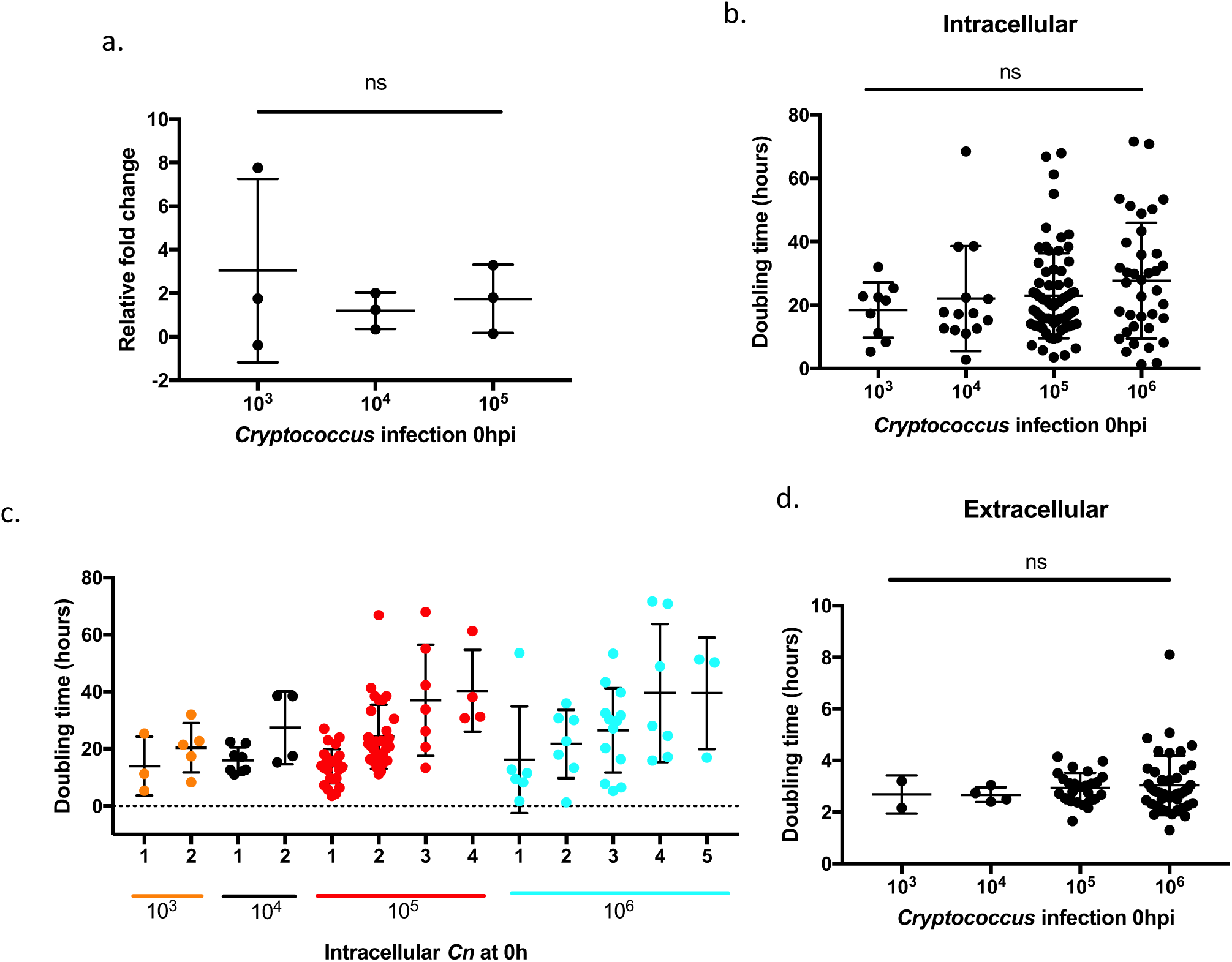
Doubling time of KN99 GFP during human MDM infection. Human monocyte derived macrophages (MDMs) were infected with an increasing number of KN99 GFP cryptococcal cells for 24 hours. The relative cryptococcal fold change was not significantly different as starting fungal burden increased (a, one-way ANOVA p = 0.696, no pairwise significance). As fungal burden increased, the doubling time of individual cryptococci which were intracellular within MDMs was not significantly different (b, one-way ANOVA p = 0.264, no pairwise significance). Within each initial fungal dose, as the number of phagocytosed cryptococci for each macrophage increased, the doubling time of the cryptococci also increased (c). When comparing doubling times within each infection burden (10^3^, 10^4^, 10^5^, 10^6^), only 10^5^ showed significantly different doubling times with increasing macrophage infection (one-way ANOVA p <0.0001). As fungal burden increased, the doubling time of individual cryptococci which are extracellular was not significantly different (d, one-way ANOVA p = 0.844, no pairwise significance). Three repeats with three replicates per repeat were performed, with up to 13 cryptococci followed per replicate. Means and standard deviations are shown.

### Comparing murine J774 macrophage and human MDM infection

To assess the similarity of the two macrophage models, we compared the infection of J774 macrophages and MDMs. For each initial fungal dose, there was no difference in the percentage of macrophages that contained cryptococci between J774 cells and MDMs at both 0h (Figure 5a Mann-Whitney test 10^3^ p > 0.999, 10^4^ p = 0.700, 10^5^ p = 0.200) and 18h (Figure 5b Mann-Whitney test 10^3^ p = 0.100, 10^4^ p = 0.400, 10^5^ p = 0.700). This suggests that J774 macrophages and human MDMs showed similar abilities to phagocytose cryptococci over increasing fungal burdens. Because fungal replication is important for propagating infection, and macrophage control of replication must be considered, we also compared the replication of cryptococci in the presence of each type of macrophage. There was no difference in intracellular cryptococcal fold change between the two models as initial fungal burden increased (Figure 5c cryptococcal fold change, Mann-Whitney test 10^3^ p = 0.700, 10^4^ p = 0.200, 10^5^ p > 0.999), though J774 macrophages replicated and MDMs did not (Figure 5d macrophage fold change, Welch’s t-test 10^3^ p = 0.0002, 10^4^ p = 0.0184, 10^5^ p = 0.0314). The doubling time of cryptococci was significantly higher for cryptococci within human MDMs (Figure 5e Mann-Whitney test 10^4^ p = 0.008, 10^5^ = 0.0002), suggesting increased control over cryptococcal proliferation rate within human MDMs. We also compared extracellular doubling time of cryptococci, though only two comparable fungal doses were considered for each assay. Extracellular proliferation was faster in the presence of human MDMs (Figure 5f Mann-Whitney test 10^4^ p = 0.0001, 10^5^ p = 0.0005). This was likely due to faster cryptococcal growth in RPMI, the media used for human MDM assays, compared to DMEM, which is used for J774 cells (data not shown; doubling time of cryptococci in SF RPMI = 3.15 hours, in SF DMEM 5.88 hours, Mann-Whitney test p < 0.0001 with at least 79 cryptococci considered for each medium). The differences in cryptococcal doubling times between the two models suggest that human MDMs enforced stronger control over intracellular replication rate.

**Figure 5.**
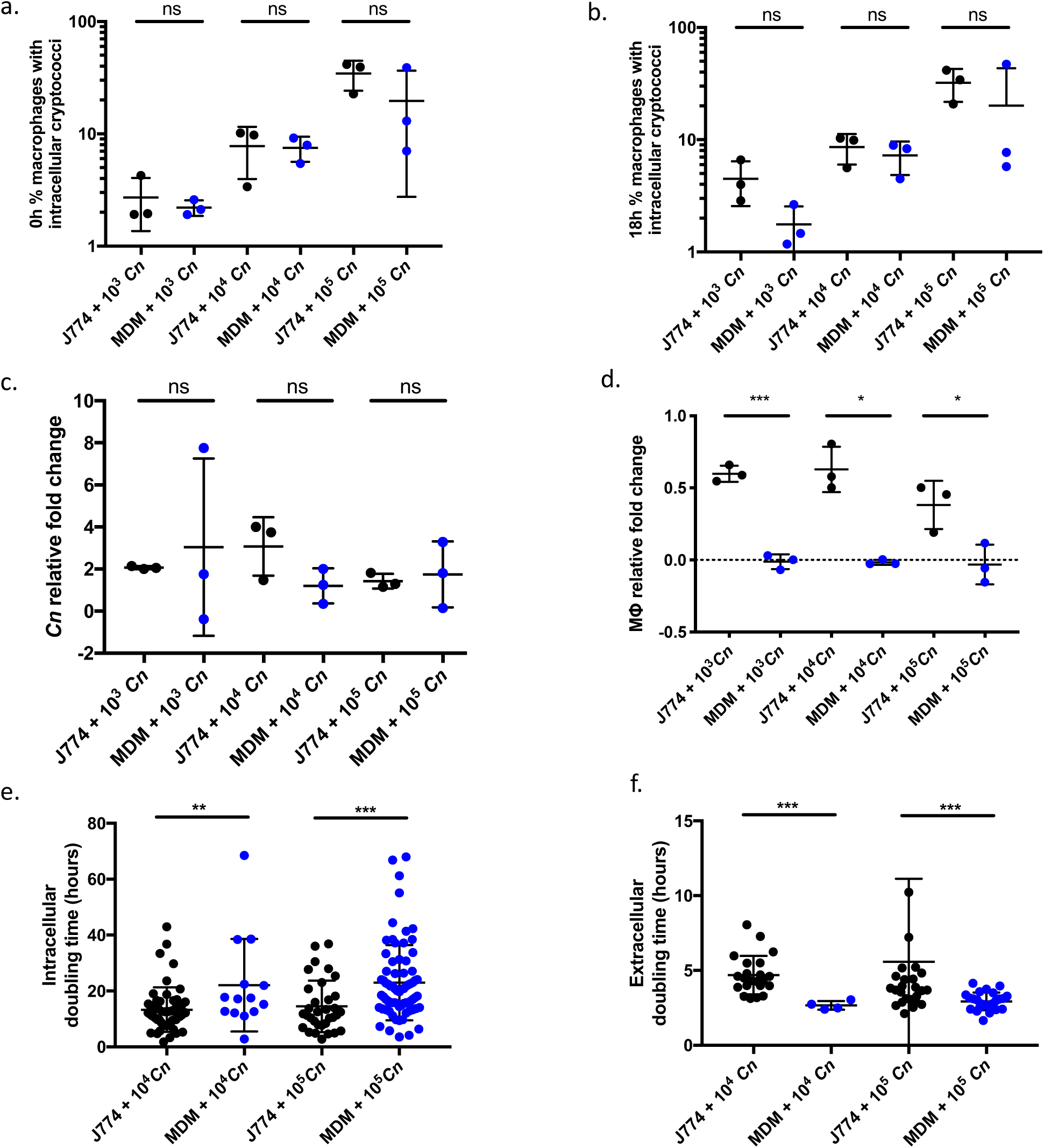
Comparing response to fungal dose by J774 macrophages and human MDMs. J774 murine macrophages (black points) and human MDMs (blue points) were infected with KN99 GFP *C. neoformans* for 18 hours. For a-d, the means of each replicate are displayed as points, and means and standard deviations shown. For e and f, points represent individual cryptococci. At 0h the percentage of macrophages which contained intracellular cryptococci over increasing initial fungal burdens was not significantly different between J774 macrophages and MDMs (a, Mann-Whitney test 10^3^ p > 0.999, 10^4^ p = 0.700, 10^5^ p = 0.200). At 18h the percentage of macrophages which contained intracellular cryptococci over increasing initial fungal burdens was not significantly different between J774 macrophages and MDMs (b, Mann-Whitney test 10^3^ p = 0.100, 10^4^ p = 0.400, 10^5^ p = 0.700). There was no significant difference in intracellular fungal fold change between J774 and MDM infection (c, Mann-Whitney test 10^3^ p = 0.700, 10^4^ p = 0.200, 10^5^ p > 0.999). While there was little change in the number of MDMs, J774 macrophages increased in number by at least 50% (d, Welch’s t-test 10^3^ p = 0.0002, 10^4^ p = 0.0184, 10^5^ p = 0.0314). Intracellular doubling time for cryptococci within MDMs was significantly higher than for cryptococci within J774 macrophages (e, Mann-Whitney test 10^4^ p = 0.008, 10^5^ = 0.0002). Doubling time for cryptococci outside of macrophages was significantly lower in the presence of human MDMs compared to J774 macrophages (f, Mann-Whitney test 10^4^ p = 0.0001, 10^5^ p = 0.0005).

## Discussion

We sought to explore how *C. neoformans* might exhibit density dependent phenotypic responses in their interactions with macrophages, as has been demonstrated in culture alone (Albuquerque et al., 2014). We hypothesised that the density of fungi would alter the interaction with macrophages, a critical step in control of cryptococcal infection (refs). However, using both mouse cell line and human primary macrophages we found no evidence of density dependent responses over a broad range of fungal cell densities. This is an important finding as it demonstrates how the change in fungal burden does not alter observed variation in outcome of infection.

During infection, many pathogens possess the ability to signal within the population once a specific population density, and signalling molecule threshold, is reached. This quorum sensing phenomenon results in coordinated gene expression and group response, and can affect virulence and pathogen interactions with host cells. Several studies have shown that *C. neoformans* may also possess quorum sensing capabilities in which genes related to population density affect fungal growth in culture (Lee et al., 2007, 2009), filamentation in unisexual mating (Tian et al., 2018), as well as melanisation and biofilm formation (Albuquerque et al., 2014). While phagocytosis and intracellular replication have been related to patient CSF burden (Sabiiti et al., 2014), no studies have closely looked at the effect of cryptococcal density on interactions between cryptococci and macrophages. Because interactions between *C. neoformans* and host macrophages are pivotal in deciding the outcome of cryptococcosis, and because increasing fungal burden during the progression of infection may affect these interactions, we investigated the effect of cryptococcal density on fungal phagocytosis and replication.

Phagocytosis is a crucial component of the immune response against pathogens, and because increasing infection burden may affect the behaviour of phagocytes, we investigated how phagocytes respond to an increasingly high multiplicity of cryptococcal infection. While some studies have suggested that quorum sensing molecules induce macrophage phagocytosis of yeasts (Vikström et al., 2005), time lapse analysis of both murine J774 macrophage and human MDM infection showed that increasing cryptococcal MOI did not affect phagocytosis by macrophages. The percentage of macrophages with intracellular cryptococcal cells did not increase or decrease disproportionately with level of infection, suggesting that macrophages neither upregulated phagocytosis, nor became overwhelmed by the larger number of cryptococci in the extracellular environment. Furthermore, because the percentage of macrophages which contained intracellular cryptococci increased linearly with burden, the relationship was likely determined by the probability of macrophages encountering cryptococci, suggesting that each macrophage-*Cryptococcus* interaction was an independent infection not regulated by cryptococcal dose or neighbouring macrophage-*Cryptococcus* encounters. This is in agreement with a previous study showing that macrophage phagocytosis of cryptococci was proportional to fungal dose in a zebrafish model of cryptococcal infection (Bojarczuk et al., 2016). These results indicate that macrophage phagocytosis of cryptococci is not affected by the level of cryptococcal infection *in vitro*.

*C. neoformans* may colonise the macrophage intracellular niche for replication and dissemination, the percentage of macrophages which successfully phagocytosed cryptococci may alternatively be seen as the proportion of macrophages which were parasitised by cryptococci (Feldmesser et al., 2000; Ma et al., 2009; Voelz et al., 2009). During the *Cryptococcus* infection here, as the initial fungal burden increased, a greater number of cryptococci came into contact with J774 macrophages and were able to successfully invade and occupy the macrophage intracellular niche. Because the relationship between burden and macrophage infection was linear, just as the macrophages did not upregulate phagocytic uptake in response to fungal dose, cryptococci did not cooperate to induce higher frequency of macrophage infiltration in response to population density – unlike *S. enterica*, which uses QS signals to express genes required for host cell invasion (Choi et al., 2012). In J774 macrophage infection, density-dependent quorum sensing between cryptococci does not seem to enhance macrophage parasitism by cryptococci, or induce an enhanced or attenuated macrophage response.

Interestingly, at the highest level of fungal infection there was a lower percentage of MDMs which successfully phagocytosed cryptococci. This was not likely due to killing of fungal cells by MDMs at the highest fungal dose, lowering the percentage of macrophages which contained live cryptococci, as fungal death was not seen during time lapse imaging. It is possible that as fungal MOI increased, MDMs became overwhelmed and failed to phagocytose cryptococci at a level proportional to dose. Alternatively, cryptococci may have cooperated at higher fungal doses to evade immune capture, or to escape from the phagocyte. This is possible, as it has been suggested that the *S. aureus agr* locus, which encodes the *Staphylococcus* quorum sensing system, is associated with endosome escape (Shompole et al., 2003). Alternatively, just as *H. capsulatum* yeasts produce cell wall factors at high density which prevent efficient phagocytosis (Rappleye et al., 2007), it is possible that cryptococci upregulated capsule production at the higher fungal doses considered here to evade phagocytosis by MDMs. However, as this was not observed during J774 macrophage infection, the most likely explanation for the reduced phagocytosis by MDMs is the high variation in uptake between donors at the highest level of fungal infection. While it is possible that the ability of MDMs to phagocytose cryptococci plateaued at higher doses, MDMs from more donors would need to be considered before accepting that no linear relationship existed between cryptococcal infection dose and uptake by MDMs. It is known that there is high donor variation in MDM control of intracellular cryptococcal proliferation (Garelnabi et al., 2018), and it stands to reason that phagocytosis may be similarly affected. Despite this, the proportion of macrophages with intracellular cryptococci did increase with fungal MOI, though the high standard deviation in donor MDMs resulted in failure to adhere to a strict linear relationship. Again, fungal burden did not strongly affect macrophage phagocytosis of yeast, indicating that this host-pathogen interaction is not affected by the level of infection *in vitro*.

Interestingly, as the number of starting intracellular cryptococci per macrophage (i.e. the number of cryptococci per phagosome) increased, the intracellular doubling time of cryptococci within J774 macrophages did not decrease. This suggests that not only did initial extracellular cryptococcal density fail to induce increased replication of cryptococci within J774 macrophages as discussed above, the intracellular population density failed to produce a density-dependent enhancement of replication as well. Additionally, as the number of cryptococci within each phagosome increased, the doubling time of the fungi did not increase, showing that the fungi did not replicate more slowly in response to intracellular population density. Here, the cryptococci did not limit growth as population density within each J774 macrophage increased. This suggests that after phagocytosis by J774 macrophages, cryptococci did not inhibit growth in order to limit consumption of nutrients and occupation of limited niche space based on cues from population density. In fact, while nutrient starvation is an important consideration, *Cryptococcus* has specific adaptations which enable survival within phagocytes during nutrient scarcity associated with intracellular infection (Fan et al., 2005; Hu et al., 2008; Johnston and May, 2010; Nolan et al., 2017; Tucker and Casadevall, 2002; Waterman et al., 2007). As such, nutrient starvation was unlikely to be a limiting factor with increasing fungal dose within macrophages during the infection time considered here. Therefore, because cryptococci did not limit replication due to nutrient limitation, cryptococcal replication was not affected by intracellular population density, suggesting that there was no effective quorum sensing occurring within each intracellular population in these conditions.

Finally, to address the relative effects of fungal dose on macrophage-cryptococcal interactions, the J774 macrophage and human MDM models were compared. While comparable in many respects, the main difference between J774 macrophage and MDM infection was the replication rate of cryptococci. In fact, the mean doubling time of intracellular cryptococci (averaged across doses) was roughly 9.6 hours longer within MDMs compared to the murine macrophages. Interestingly, extracellular doubling time was faster in the presence of MDMs, though this was due to the different media. Nonetheless, cryptococcal replication was much slower within human MDMs, suggesting that these cells more effectively suppressed the cryptococcal growth rate. However, caution is warranted when drawing conclusions by comparing different models, as human MDMs and murine J774 macrophages do not necessarily exhibit identical activation states (Lacey et al., 2012; Levitz and Farrell, 1990). It is also important to consider that the human MDMs may have shown bias in polarisation; human MDMs are typically classified as being M2-polarised (Bazzi et al., 2017), which is permissive to intracellular cryptococcal growth – though this potential polarisation still inhibited cryptococcal replication compared to J774 macrophages. It is also possible for the macrophages to shift their polarisation state, especially after introduction of infection or cytokines (Davis et al., 2013; Lacey et al., 2012), a phenomenon that may be interesting to dissect in the future. Here, we showed that J774 macrophages and human MDMs respond similarly to increasing cryptococcal burden, though human MDMs more effectively slow the intracellular proliferation rate of *C. neoformans*.

The results discussed here have shown that increasing cryptococcal infection does not affect macrophage-*Cryptococcus* interactions *in vitro*, and that QS is not likely to be utilized by cryptococci to regulate proliferation or macrophage parasitism during J774 and MDM infections. Furthermore, because phagocytosis increased with MOI, each macrophage-*Cryptococcus* encounter may be viewed as an independent infection. Collectively, these results illustrate that macrophage-*Cryptococcus* interactions are maintained over an increasing MOI *in vitro*, that fungal density and QS responses are not likely to affect encounters between macrophages and cryptococci, and that fungal burden is unlikely to result in increased or decreased macrophage control over infection at the cryptococcal doses considered *in vitro* here.

## Experimental procedures

### *Cryptococcus* strains, growth conditions, and opsonisation

*Cryptococcus neoformans* var. grubii, strain KN99 GFP, was used for all experiments described. *Cryptococcus* was grown for 18 hours overnight in 2 ml yeast peptone dextrose (YPD 50 g/L Fisher, 2% agar Sigma-Aldrich) medium. 1 ml of *Cryptococcus neoformans* suspension was removed from the overnight preparation, and yeasts were washed and pelleted in phosphate buffered saline (PBS) using a centrifuge (3300g for 1 minute) three times. The number of cryptococci per ml was determined, and cryptococci were diluted to the required concentration. Cryptococci were opsonised using 0.1 μl of 18B7 (a monoclonal antibody to cryptococcal capsule) per 100 ml, rotating at room temperature for one hour.

### Murine macrophage culture and preparation

J774A.1 murine macrophages were maintained in DMEM (Sigma-Aldrich) containing 10% fetal bovine serum, 1% L-glutamine (Sigma-Aldrich), and 1% penicillin/streptomycin antibiotics (Sigma-Aldrich). Cells were maintained at 37°C and 5% CO_2_. Cells were used only during passages 3 through 14. Macrophages were plated in 24-well plates at 10^5^ cells per well 18 hours before experimentation. During experimentation macrophages were maintained in DMEM without fetal bovine serum, and were activated using 15 μl of 0.01 mg/ml phorbol myristate acetate (PMA) for 45 minutes before infection.

### Human cell ethics, isolation, and maintenance

Informed written consent was obtained before whole blood was collected from healthy human donors. The South Sheffield Research Ethics Committee (REC) provided ethical approval for the collection of blood for the study of monocyte derived macrophages (REC reference 07/Q2305/7).

After collection of blood samples, whole blood was added to Ficoll-Paque in a sterile centrifuge tube at a ratio of 1:2 (Ficoll:Blood). This was centrifuged at 1500 rpm for 23 minutes before the plasma layer was discarded. PBS was added to PBMCs, and this was centrifuged at 1000 rpm at 4°C for 13 minutes. Supernatant was removed, pellet was re-suspended in PBS, and was centrifuged again. Pellet was then resuspended in complete RPMI (see appendix) and 1 ml was added per well in a 24 well plate, or 3 ml was added to individual glass-bottomed dishes.

MDMs were maintained in RPMI containing 10% fetal bovine serum, 1% L-glutamine (Sigma-Aldrich), and 1% penicillin/streptomycin antibiotics (Sigma-Aldrich). Macrophages were used two weeks after isolation to allow macrophage differentiation from monocytes. During experimentation MDMs were kept in serum free media, and no activation was performed.

### Macrophage infection

Macrophages were infected with *Cryptococcus* and incubated at 37°C, 5 % CO_2_ for two hours. Extracellular cryptococci were then removed by washing the macrophages three times in 37°C PBS before being resuspended in serum free media.

### Time lapse imaging and analysis

For both J774 and MDM imaging, Nikon eClipse TI microscope and NIS Elements 4.51 analysis software were used. x20 phase imaging was performed using perfect focus to maintain clear images throughout the duration of imaging. Differential interference contrast (DIC) images were captured every 2-3 minutes, and GFP images were captured every 30 minutes to follow intracellular cryptococci. Phagocytosis and replication quantifications were performed by manually watching individual cryptococci throughout the course of the time lapse assays and counting events manually.

The percentage of macrophages which contained intracellular cryptococci was determined by counting all of the macrophages in a field of view and determining which contained intracellular cryptococci using the GFP channel. The percentage of infected macrophages per field of view was converted to macrophages per ml based on known size of the field of view and the 24 well plates. Intracellular replication of cryptococci was determined by following individual cryptococci using the GFP channel and noting when a fungal cell budded. Extracellular proliferation was quantified in the same way. The number of cryptococci at different times was plotted in GraphPad Prism (versions 7.0c and 8.2.1) and the doubling time was determined by plotting each using an exponential fit.

### Flow cytometry performance and analysis

After either 0 or 18 hours of infection, extracellular cryptococci were washed from the wells three times using warmed PBS. 200 μl of accutase was added for 5 minutes to dislodge macrophages from the plate before the addition of 800 μl of PBS and collection of cells into Eppendorf tubes. Flow cytometry was performed using Attune™ Nxt acoustic focusing cytometer (Life Technologies and Attune™ NxT Software v2.5). Samples were mixed by hand, and 200 μl was run through the flow cytometer. The macrophage population was determined by known size and complexity characteristics. Macrophages with increased GFP fluorescence (BL1A) were determined to be those containing KN99 GFP *Cryptococcus neoformans*, and the percentage of macrophages containing this enhanced GFP signal were calculated as a proportion of the total number of macrophages detected.

### Statistics

GraphPad Prism versions 7.0c and 8.2.1 were used for statistical analyses. Normality was determined based on adherence to a Gaussian distribution. Normality tests were not performed. 95% confidence intervals were used to assess statistical significance. Unpaired two-tailed t-tests were performed to compare the means of conditions. Mann-Whitney tests were performed when data was non-parametric. Ordinary one-way ANOVAs were performed to compare the means of conditions. Multiple comparisons were performed using Tukey’s test to correct for multiple comparisons.

---

**Supplemental Fig. 1.**
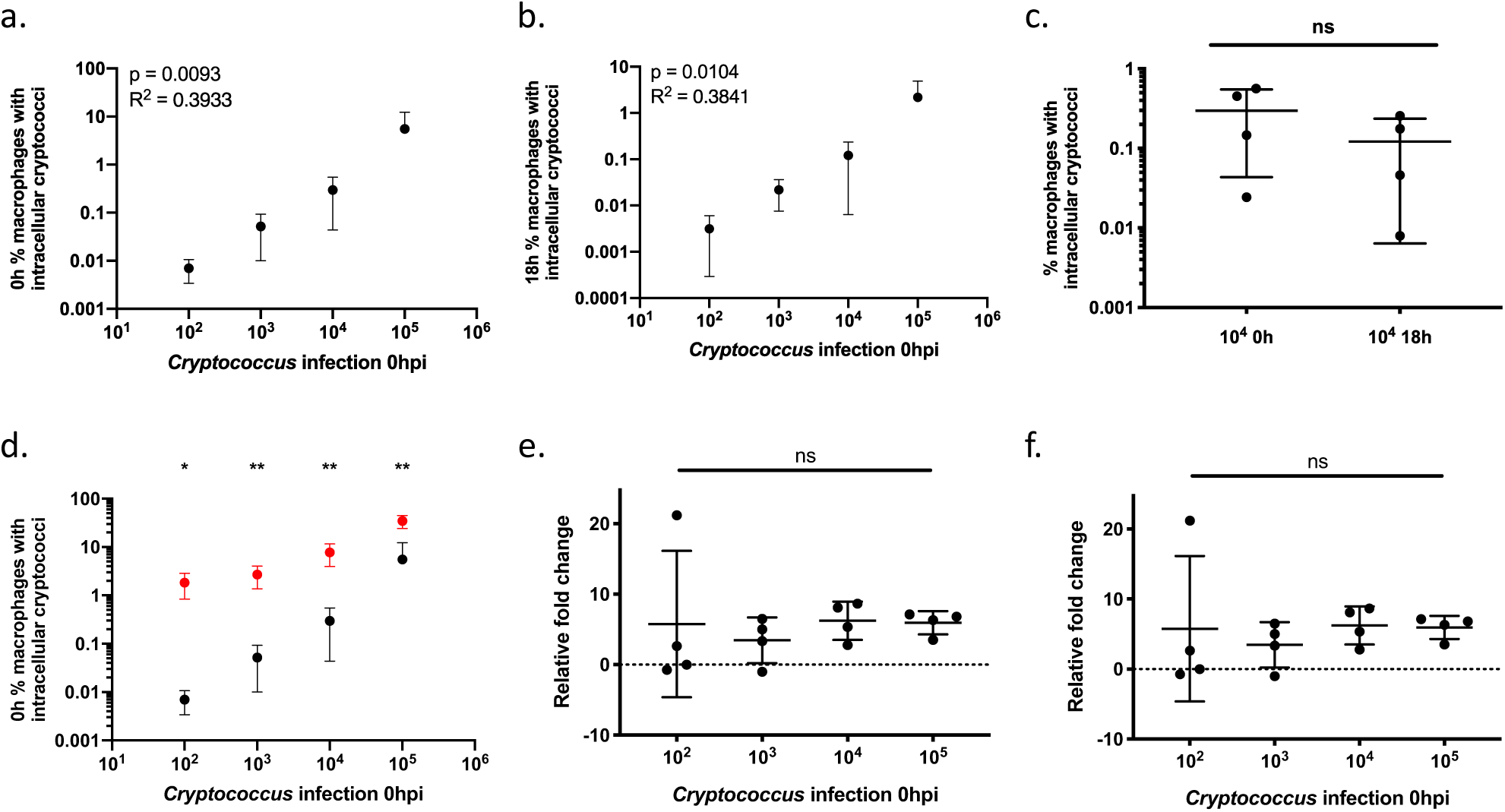
Flow cytometry analysis of J774 murine infection. J774 macrophages were infected with KN99 GFP *C. neoformans* and infection was quantified by flow cytometry. At 0h the percentage of macrophages which contained intracellular cryptococci was linearly related to fungal burden (a, linear regression p = 0.0093, R^2^ = 0.393). At 18h the percentage of macrophages which contained intracellular cryptococci linearly related to fungal burden (b, linear regression p = 0.0104, R^2^ = 0.3841). The proportion of macrophages containing cryptococci decreased but was not significantly different over 18 hours (c, Mann-Whitney test p = 0.4857); only one initial fungal dose is shown here for the sake of clarity, but results are similar for each comparison. The percentage of macrophages at 0h which contained intracellular cryptococci increased linearly with increasing MOI (d, linear regression flow cytometry p = 0.0009, R^2^ = 0.9981: time lapse p = 0.0030, R^2^ = 0.994). Similar results were achieved for 18h (not shown, linear regression flow cytometry p = 0.0008, R^2^ = 0.9984; linear regression timelapse p = 0.0016, R^2^ = 0.9969). The percentage of infected macrophages calculated by timelapse analysis was significantly higher than when calculated by flow cytometry (unpaired t-test 10^2^ p = 0.0130, 10^3^ p = 0.0095, 10^4^ p = 0.0098, 10^5^ p = 0.0061). The relative macrophage fold change was not affected by increasing fungal dose (e, one-way ANOVA p = 0.9980). There was no significant difference in intracellular fungal fold change over increasing levels of initial cryptococcal infection (f, one-way ANOVA p = 0.8922). Four repeats were performed, and the average of 6 replicates per condition is shown here. Means and standard deviations are shown. * p < 0.05, ** p < 0.01.

